# Intestinal epithelial and intraepithelial T cell crosstalk mediates a dynamic response to infection

**DOI:** 10.1101/154237

**Authors:** David P. Hoytema van Konijnenburg, Bernardo S. Reis, Virginia Pedicord, Julia Farache, Gabriel D. Victora, Daniel Mucida

## Abstract

Intestinal intraepithelial lymphocytes (IELs) are located at the critical interface between the intestinal lumen, which is chronically exposed to food and microbes, and the core of the body. Using high-resolution microscopy techniques and intersectional genetic tools, we investigated the nature of IEL responses to luminal microbes. We observed that TCRγδ IELs exhibit distinct location and movement patterns in the epithelial compartment that were microbiota-dependent and quickly altered upon enteric infections. These infection-induced changes included increased inter-epithelial cell (EC) scanning, anti-microbial gene expression and glycolysis. Direct modulation of glycolysis was sufficient to change γδ IEL behavior and susceptibility to early pathogen invasion. Both γδ IEL behavioral and metabolic changes were dependent on EC pathogen sensing. Our results uncover a coordinated EC–IEL response to enteric infections that modulates lymphocyte energy utilization and dynamics and supports maintenance of the intestinal epithelial barrier.

## Introduction

In order to function properly, the immune system depends on the capacity of its cells to surveil compartments of the body and respond to particular environmental cues. Intestinal intraepithelial lymphocytes (IELs) comprise one of the most abundant T cell populations and potentially provide a first line of immune defense against pathogens via their location at the critical interface between the intestinal lumen and the core of the body (Ismail et al., 2009; Jabri and Abadie, 2015). IELs constitute a heterogeneous group of “activated yet resting” T lymphocytes characterized by high expression levels of activation markers, gut-homing integrins, NK-like receptors, cytotoxic T lymphocyte (CTL)-related genes and anti-inflammatory or inhibitory receptors (Cheroutre et al., 2011; Denning et al., 2007). The main populations of “natural” IELs are TCRγδ^+^CD8αα^+^ and TCRαβ^+^CD8αα^+^ cells (Guy-Grand et al., 1991; Huang et al., 2011; Lefrancois, 1991). Additionally, peripheral, mature TCRαβ CD4^+^ and CD8^+^ T cells can acquire IEL markers upon migration to the intestine (“peripheral” IELs) (Das et al., 2003; Mucida et al., 2013; Reis et al., 2013).

Irrespective of subtype, tightly regulated control of IEL function is crucial for the maintenance of the epithelial cell barrier and gut physiological inflammation (Tang et al., 2009). Inappropriate activation of the CTL capacity of IELs can induce chronic inflammatory disorders such as celiac disease and IBD (Meresse et al., 2006; Tang et al., 2009), while a lack of IELs leads to impaired protection against bacterial infection (Ismail et al., 2009; Ismail et al., 2011). Although marker-based studies underscore several features related to the phenotype, development and migration of IELs (Klose et al., 2014; Li et al., 2011; Malamut et al., 2010; McDonald et al., 2014; Meresse et al., 2004; Meresse et al., 2006; Reis et al., 2014; Sujino et al., 2016), dissecting the role played by these cells under homeostatic and pathophysiological conditions has been hampered by their poor survival in culture and difficulty assessing IEL function *in vivo* (Cheroutre et al., 2011; Edelblum et al., 2012; Edelblum et al., 2015; Vantourout and Hayday, 2013).

To investigate how IELs respond to enteric microbes, we established intravital and 3D deep-tissue imaging tools in mouse models of infection or microbiota manipulation. We found that γδ IELs are predominantly located in the middle and upper region of intestinal villi, are highly motile, and display a structured migration pattern, suggesting an epithelial surveillance program. Pathogenic bacterial or protozoan infections induced a dynamic response by γδ IELs, rapidly changing their motility and pattern of movement between intestinal epithelial cells (ECs). This swift γδ IEL response was associated with enhanced anti-microbial gene expression and a metabolic switch towards glycolysis, even in the absence of active proliferation. Pharmacological modulation of energy utilization pathways recapitulated infection-induced changes and resulted in altered susceptibility to early pathogen invasion. Using inducible, EC-specific *Myd88* conditional knockout mice, we demonstrated that the γδ IEL metabolic switch, movement behavior and immune response required pathogen sensing by surrounding ECs. Our results uncover a coordinated EC–IEL response to luminal microbes that modulates lymphocyte energy utilization and cell dynamics within the epithelium, thus supporting the maintenance of epithelial barrier defense.

## Results

### γδ IELs display microbe-dependent motility patterns

The gut epithelial compartment represents a spectrum of increasingly differentiated intestinal epithelial cells (ECs) from crypt to villus tip, which corresponds to increasing levels of microbe exposure (Ismail et al., 2011; Sato et al., 2009). Likewise, microbial content and diversity increases from proximal (duodenum) to distal (ileum) small intestine and to large intestine (Donaldson et al., 2016). Additionally, the number and distribution of IEL subpopulations also significantly changes along the intestine. While germ-free (GF) mice show drastic reductions in “peripheral IEL” numbers and frequency along the intestine when compared to their conventionally-housed, specific pathogen-free (SPF) counterparts (Bandeira et al., 1990; Sujino et al., 2016), overall numbers of “natural IELs” remain mostly intact, particularly γδ IELs (Fig. S1A). To define γδ IEL localization within the intact epithelial compartment, we performed tissue clearing using iDISCO (Renier et al., 2016) and *FocusClear*™ (which provided similar results, Movie S1A, B) and subsequent deep-tissue 3D imaging of intact small intestine segments from TCRγδ^GFP^ reporter mice using light-sheet, confocal and multi-photon microscopy (Gabanyi et al., 2016). Resulting images and automated quantification revealed that the majority of γδ IELs accumulate in the middle of the villus and fewer cells in the tip or bottom of the villus, including the crypts (Movie S1A, B; Figure 1A, B, Figure S1B, C). To address how luminal microbes influence γδ IEL localization, we re-derived TCRγδ^GFP^ reporter mice into germ-free (GF) conditions. A significant shift towards the crypts was observed for γδ IELs in the duodenum, jejunum and ileum of GF animals. In contrast to SPF mice, we did not observe γδ IEL accumulation in specific regions of the villi (Figure S1B, C; Figure 1A, B). Conventionalization of GF TCRγδ^GFP^ mice with microbiota from SPF TCRγδ^GFP^ mice rescued IEL positioning in the ileum, while broad-spectrum antibiotic treatment of SPF TCRγδ^GFP^ mice mirrored GF IEL positioning (Figure 1C).

**Figure 1.**
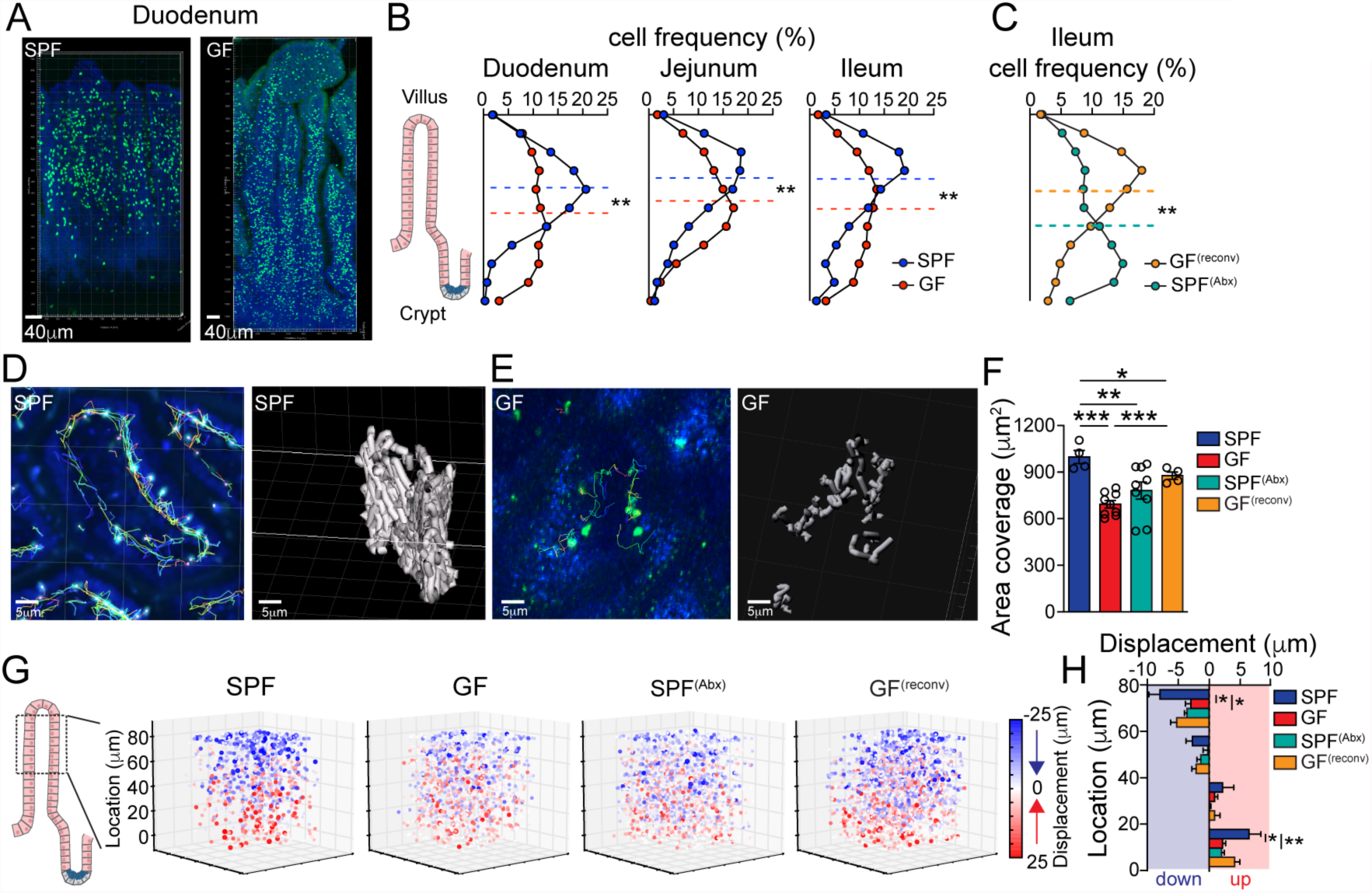
Steady-state behavior of intestinal IELs. (A) Representative image of cleared (*FocusClear*™) duodenum villi of SPF and GF TCRγδ^GFP^ mice (see Movies S1A, 1B). In green (GFP), TCRγδ^+^ cells and in blue (Hoechst), EC nuclei. (B) Frequency of TCRγδ^+^ cell distribution along the duodenum, jejunum and ileum villi of SPF or GF mice. Dashed line indicates mean of median positions for 4 mice/group. ** = p<0.01 with two-tailed Students *t* test. (C) Frequency of TCRγδ^+^ cell distribution along the ileum villi of 4 weeks broad-spectrum antibiotics – treated SPF mice (SPF^ABX^) or GF mice, re-conventionalized (7 days) with SPF microbiota (GF^reconv^). Dashed line indicates mean of median positions for 4 mice/group. ** = p<0.01 with two-tailed Students *t* test. (D-H) Intravital microscopy (IVM) analyses of TCRγδ^GFP^ SPF mice or SPF mice treated for 4 weeks with antibiotic mix (SPF^ABX^) prior to analysis, and GF mice or GF mice colonized with SPF microbiota (GF^reconv^) 7 days prior to analysis (see Movies S2A-D). (D, E) Tracking of TCRγδ^GFP^ cells (colorful lines, left panels) in 3D over time was performed using Imaris. (D, E) 3D reconstruction (grey cylinders, right panels) of the area in a villus covered by TCRγδ^GFP^ cells in 30 mins. (F) Quantification of unique area covered/IEL/hour. Means and SEM shown, each dot = 1 movie, n = at least 4 mice/group in 3 independent experiments. * = p<0.05, ** = p<0.01, *** = p<0.001 with two-tailed Students *t* test.(G, H) Visualization (G) and quantification (H) of TCRγδ vertical (*Z*) displacement. Pooled imaging data from SPF, GF, SPF^ABX^ and GF^reconv^ mice is shown. Panels show starting position and mean vertical displacement over time for each individual cell within each movie. Color density and size indicate degree of *Z* displacement down- (blue) or upwards (red). Graph shows mean *Z* displacement per anatomical villus region as indicated.

We next used deep (+/- 80 μm) multi-photon intra-vital microscopy (IVM) to gain insight into the cell dynamics associated with γδ IEL distribution. Previous reports using time-lapse live imaging of superficial 15 μm villus sections revealed that γδ IELs are highly dynamic and able to respond to directly applied pathogenic bacteria (Edelblum et al., 2012; Edelblum et al., 2015). Our live 4D imaging and supervised quantification showed that under homeostatic conditions γδ IELs migrate almost exclusively in the space between the epithelial layer and the basement membrane, rapidly covering the entirety of the villous epithelium (Movie S2A). We found that in naïve SPF, a single γδ IEL covered about 950 μm^2^ of unique basolateral epithelial surface area (or ~2.2% of a villus) per hour. In contrast, each γδ IEL from GF or SPF mice treated with broad-spectrum antibiotics covered only roughly 650 μm^2^ of unique area (or 1.5% of the villus epithelium) per hour (Movie S2B, C; Figure 1D-F), despite the overall maintenance of a cell speed of ~6 μm/min (Figure S1D). Furthermore, at one-week post conventionalization of GF mice we observed a partial recovery of γδ IEL area coverage (Movie S2D; Figure 1F). Because our cleared tissue analyses indicated that γδ IEL positioning within the villi is influenced by the microbiota, we quantified their vertical displacement using IVM. IELs occupying the tips of villi or the regions closer to the crypt showed a strong vertical migration pattern (Figure 1G, H). Nevertheless, the γδ IEL downward displacement exceeded the upward, with net displacement for γδ IELs around 4-6 μm/hour towards the crypts. GF mice showed a significantly altered γδ IEL vertical displacement and reduced total net displacement to roughly 0.5 μm/hour (Figure 1G, H). Conventionalization of GF mice rescued γδ IEL vertical displacement, while four-week treatment of SPF mice with broad-spectrum antibiotics mirrored GF γδ IEL dynamics (Figure 1G, H). These results suggest that population migration dynamics of non-renewing γδ IELs may represent an active mechanism for their retention in the rapidly renewing epithelial layer that is modulated by exposure to commensal microbiota. Together, the above data indicate that the motility of γδ IELs along the epithelial layer is directed by commensal bacteria, suggesting an organized surveillance behavior able to cover the entire epithelial surface within hours.

### Infection induces rapid behavioral changes in TCRγδ^+^ IELs

The observation of a microbiota-dependent IEL epithelial motility pattern in the steady state led us to speculate about a possible role for this behavior in responses to enteric infections, previously suggested by studies using TCRγδ-deficient mice (Ismail et al., 2011; Li et al., 2012) and time-lapse imaging of intestine bathed in medium containing *Salmonella* (Edelblum et al., 2015; Ismail et al., 2011; Li et al., 2012). We first analyzed γδ IEL behavior after oral infection with either wild-type *Salmonella enterica* Typhimurium or a mutant *invA* strain with reduced pathogenicity due to its impaired invasion capacity (Galan et al., 1992). While γδ T cells in both naïve and infected mice maintained their presence almost exclusively in the epithelial compartment, the motility pattern of these cells changed sharply upon infection. As early as two hours post-infection with *Salmonella*, γδ IEL covered a reduced area and their movement became serpentine with significantly increased movement between ECs and into the lateral intercellular spaces, a behavior we denominated “flossing”. This flossing movement doubled upon infection despite the overall maintenance of a cell speed of ~6 μm/min (Movie S3A; Figure S2A, Figure 2A-D). This pattern persisted for at least 18 hours in mice infected with wild-type *Salmonella*, but was significantly reduced in mice infected with the *invA* mutant, suggesting that invasion or persistence of the pathogen is required to sustain this behavior (Movie S3B; Figure 2A-D). While flossing occurred sporadically and at a lower frequency (around 35 flosses/100 cells) in naïve mice, regions with a high frequency of flossing (“hotspots”) were observed in *Salmonella*-infected animals, particularly at lower regions of the villi (Figure S2B; Figure 2E). Additionally, γδ IELs from wild-type *Salmonella*-infected mice showed reduced vertical displacement with an increased shift of γδ IEL distribution towards the bottom parts of villi (Figure S2C; Figure 2F, G). These analyses indicate that *Salmonella* infection swiftly induces a set of motility changes in γδ IELs that is distinct from the one observed in response to commensal microbiota.

**Figure 2.**
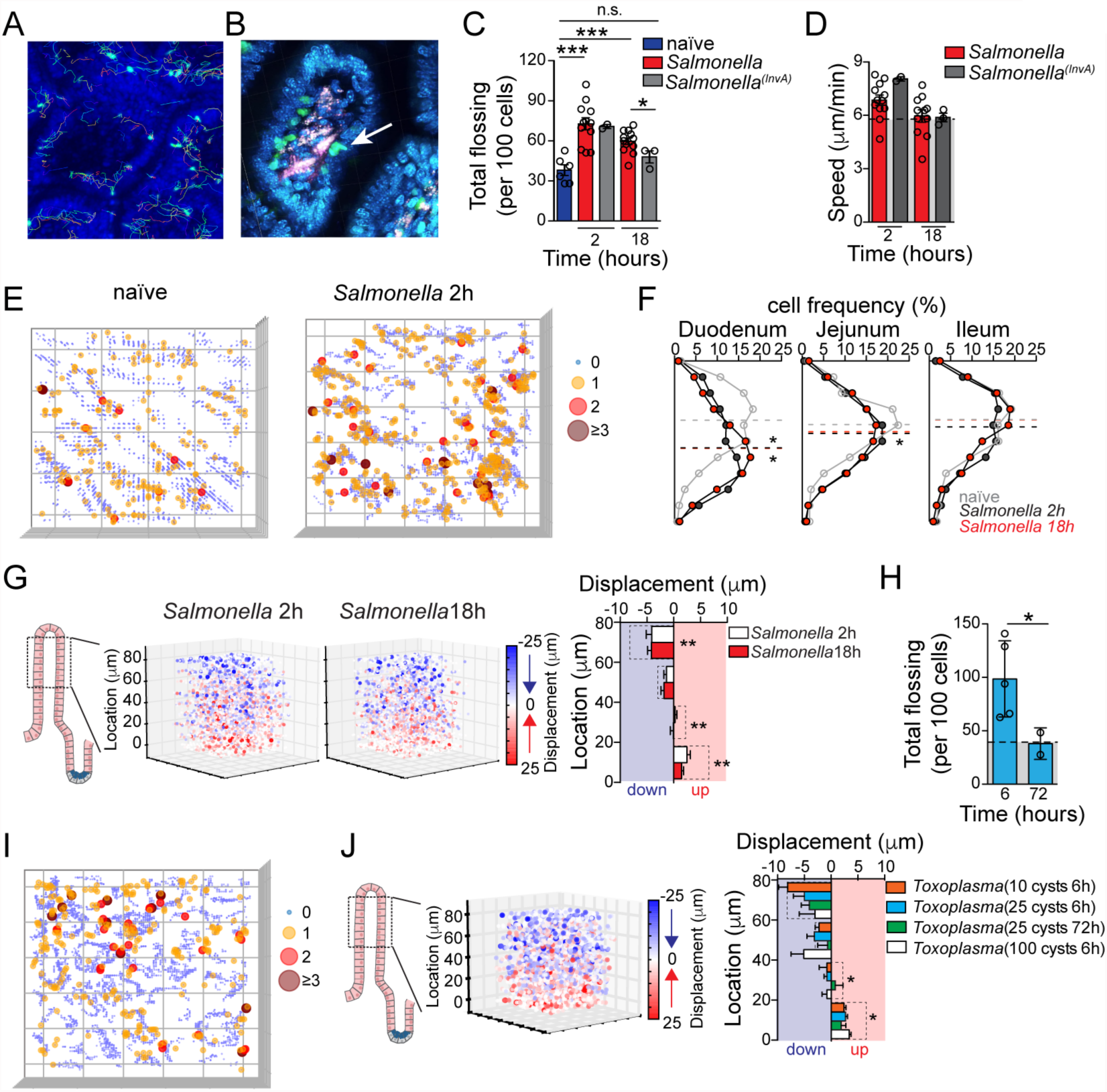
Influence of enteric infections on IEL behavior. (A-G) IVM analyses of TCRγδ^GFP^ mice post *S.* Typhimurium (or *invA* mutant) infection. (A) Tracking of TCRγδ^GFP^ cells (colorful lines) in 3D over time was performed using Imaris. (B) Zoom of 1 villus showing a flossing (arrow) movement. (C) Unbiased computational quantification of inter-epithelial cell movements (flossing) (see Movies S3A-C). Each dot = 1 movie, n = at least 4 mice/group in 3 independent experiments. * = p<0.05, *** = p<0.001 with two-tailed Students *t* test. (D) Means and SEM of TCRγδ^GFP^ cell speed are shown for indicated infections and timepoints, each dot = 1 movie. (E) Representative visualization of frequency of overlapping flossing events (“hotspots”) at unique coordinates, ranging from 0 (grey) to 3 or more (maroon). Luminal to crypt orientation of a 3D plot is shown. (F) Frequency of TCRγδ^GFP^ cell distribution along cleared duodenum, jejunum and ileum villi of 2h or 18h *S.* Typhimurium*-*infected and naïve TCRγδ^GFP^ mice. Dashed line indicates mean of median positions for 4 mice/group. * = p<0.05 with two-tailed Students *t* test. (G) IVM visualization and quantification of TCRγδ vertical (*Z*) displacement. Panels show starting position and mean vertical displacement over time for each individual cell within each movie. Color density and size indicate degree of *Z* displacement down- (blue) or upwards (red). Graph shows mean *Z* displacement per anatomical villus region as indicated. Dashed line indicates SPF naïve values (Fig. 1). N = at least 4 mice / group in min. 3 independent experiments. ** = p<0.01 (versus SPF naïve) with two-tailed Students *t* test. (H) Unbiased computational quantification of flossing movements after infection with 25 cysts of *Toxoplasma gondii* visualized by IVM (see Movie S3C). * = p<0.05 (vs. SPF naïve, from panel 2C) with two-tailed Students *t* test. Toxoplasma: each dot = 1 mouse, SPF naïve: Each dot = 1 movie, n = at least 4 mice/group in 3 independent experiments. (I) Representative visualization of frequency of flossing hotspots events as in “D”. (J) Visualization and quantification of TCRγδ vertical (*Z*) displacement. Pooled imaging data from mice at indicated time points (as in “F, G”) after *Toxoplasma* infection with indicated number of cysts is shown. Dashed line indicates SPF naïve values (Fig. 1). N= at least 4 mice/group 3 independent experiments. * = p<0.05 (versus SPF naïve) with two-tailed Students *t* test.

To evaluate whether this γδ IEL behavioral response to *Salmonella* could be extended to additional small intestine pathogens, we also investigated IEL movement following *Toxoplasma gondii* infection, another pathogen to which IEL function has been previously linked (Cohen and Denkers, 2014; Dalton et al., 2006; Edelblum et al., 2015; Li et al., 2012). We found that infection with as few as 25 *Toxoplasma* cysts drastically altered γδ IEL behavior in a manner similar to that observed during wild-type *Salmonella* infection, inducing roughly 100 flosses/100 cells per hour, preferentially at hotpots (Movie S3C; Figure S2B, D; Figure 2H, I). However, hotspots after *Toxoplasma* infection were observed closer to tip regions of the villi (Figure S2B). *Toxoplasma* infection also induced dose-dependent changes in γδ IEL vertical displacement, leading to a significant increase in IELs that remained at the same height in the villi as they started (Figure 2J). γδ IEL movement patterns returned to normal by 72 h post-*Toxoplasma* infection, a time-point at which the initial invasion by tachyzoites through the epithelial layer has been completed (Barragan and Sibley, 2002)(Figure 2H). These rapid but temporary changes in IEL behavior, also observed after *invA Salmonella* infection, suggest that γδ IELs are exquisitely responsive to the nature and continued presence of pathogens. Together, the above imaging analyses suggest γδ IEL programs for steady state surveillance and a swift reaction to luminal pathogens, characterized by specific cell dynamics and positioning in the tissue.

### Coordinated EC-IEL responses to enteric pathogens

To gain insights into the mechanisms responsible for γδ IEL behavioral responses to luminal microbes and the role of surrounding ECs in this process, we performed parallel transcriptome analyses on sorted γδ IELs and ECs from wild-type *Salmonella-*infected and naïve control mice. *Salmonella* infection resulted in increased γδ T cell expression of genes associated with bacterial defense responses, particularly activating and inhibitory cytotoxic (NK-like) receptors, as well as adhesion molecule pathways and cytoskeletal rearrangement-associated genes. Infection also induced changes in immune response-related pathways in sorted ECs, including TLR sensing and downstream Myd88 signaling (Figure S3A, B and Figure 3A, B). Moreover, in both γδ T cells and ECs we observed an upregulation of the Wnt/β-Catenin pathway upon infection, a pathway usually associated with changes in the epithelial cell turnover rate and tissue regeneration (Clevers et al., 2014; Karin and Clevers, 2016) and cellular metabolism (Inoki et al., 2006). The activation of microbe-sensing pathways by ECs, associated with similar gene expression changes in both ECs and IELs in immune response-related and metabolic pathways, pointed to ECs as potential primary microbe-responding cells that could prompt neighboring IELs.

**Figure 3.**
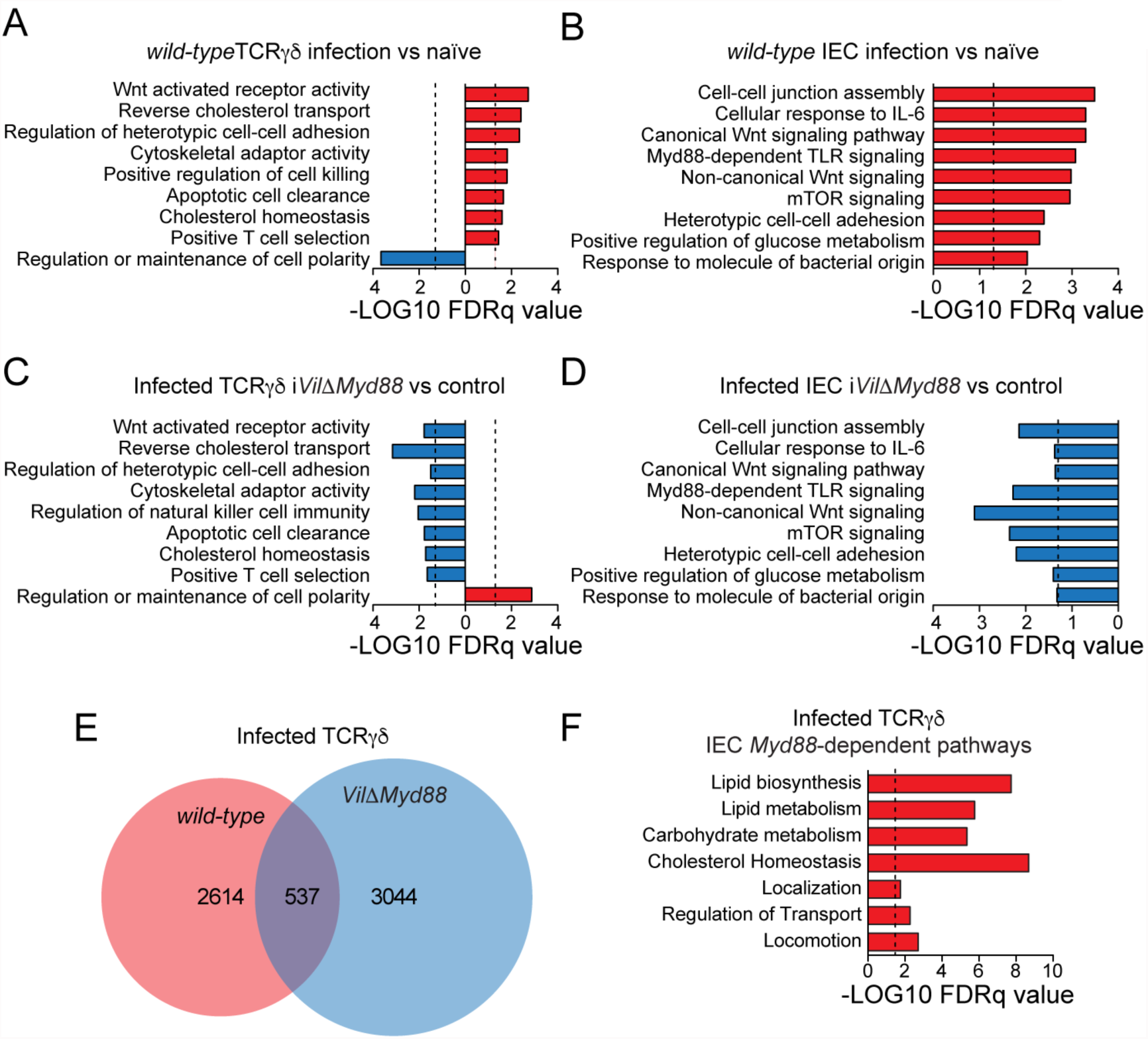
Myd88 expression by ECs is required for a coordinated epithelial transcriptional response to microbes. Sorted TCRγδ IELs (CD45^+^EpCAM^-^TCRαβ^-^TCRγδ^+^CD8α^+^) and ECs (CD45^-^EpCAM^+^) were isolated from *S.* Typhimurium-infected or naïve wild-type mice (A, B) or from tamoxifen-treated *S.* Typhimurium-infected i*Vil*Δ*Myd88* and Cre^−^ littermate control mice (C, D). RNA-sequencing was performed and Gene Set Enrichment Analysis (GSEA PreRanked, Broad Institute) used to identify GO Biological Processes enriched in TCRγδ IELs (A, C) or ECs (B, D). Expression of individual genes is shown in Figure S3. (E) Top genes upregulated in wildtype TCRγδ cells upon infection (red) were cross-referenced with the top genes downregulated in TCRγδ cells from infected i*Vil*Δ*Myd88* mice (blue). (F) For genes shown to overlap in (E), enrichment analysis was performed using the GO/PANTHER database overrepresentation test. N = 3 animals per group for wildtype and TCRγδ and EC as well as i*Vil*Δ*Myd88* and control EC. N = 2 animals per group for i*Vil*Δ*Myd88* and control TCRγδ.

To investigate whether EC microbe sensing played a role in the coordinated EC-IEL response to infection, we analyzed *Villin*^CreER^*Myd88*^f/f^ (i*Vil*Δ*Myd88*) mice, which feature tamoxifen-inducible ablation of the adaptor protein Myd88 specifically in ECs (Pedicord et al., 2016). Similar to analyses performed in wild-type mice, we evaluated the transcriptomes of sorted ECs and γδ IELs in tamoxifen-treated i*Vil*Δ*Myd88* mice after infection (Figure S3A, B; Figure 3C, 3D). As expected, upregulation of Myd88-dependent immune response genes was abrogated in ECs isolated from i*Vil*Δ*Myd88* mice post-infection. A similar impairment was observed for the mTOR and β-Catenin/Wnt signaling pathways (Figure 3D). Although a TLR-independent, IL1Rα-inflammasome role for Myd88 signaling has been described (Brown et al., 2013), we were unable to detect significant upregulation of genes associated with this pathway or of IL-1β protein in sorted ECs from i*Vil*Δ*Myd88* or littermate control mice after *Salmonella* infection (Figure S3C). These data indicate that microbe sensing via TLRs and downstream signaling through Myd88 is the main driver for these effects in ECs 18 h post-infection. Correspondingly, γδ IELs isolated from *Salmonella-*infected tamoxifen-treated i*Vil*Δ*Myd88* mice displayed complete or partial abrogation of the transcriptional changes observed upon *Salmonella* infection in wild type mice or tamoxifen-treated Cre^-^ control mice (Figure 3C). Further analysis of the top genes upregulated in γδ IELs upon *Salmonella* infection in an EC-specific Myd88-dependent manner identified genes associated with T-cell metabolism, motility and migration (Figure 3E, F). These results suggest that key gene pathways induced in γδ IELs early after infection, particularly metabolic and motility pathways depend on microbe sensing by neighboring ECs via Myd88.

To investigate whether EC-specific Myd88 expression could directly regulate γδ IEL movement behavior in the steady state, we crossed i*Vil*Δ*Myd88* with TCRγδ^GFP^ mice (i*Vil*Δ*Myd88-*TCRγδ^GFP^). At one week after tamoxifen treatment, sufficient time for replacement of entire villi with *Myd88*-deficient ECs, we did not observe altered γδ IEL positioning or behavior between i*Vil*Δ*Myd88-*TCRγδ^GFP^ and littermate control mice (Movie S4A, Figure 4A-C). However, four weeks after tamoxifen treatment, γδ IEL dynamics acquired a “GF-like” profile, with similar vertical distribution (Figure 4A), vertical movement dynamics (Figure 4B) and unique area coverage (Movie S4B, Figure 4C), suggesting that EC sensing of the microbiota impacts IEL behavior in the steady state.

**Figure 4.**
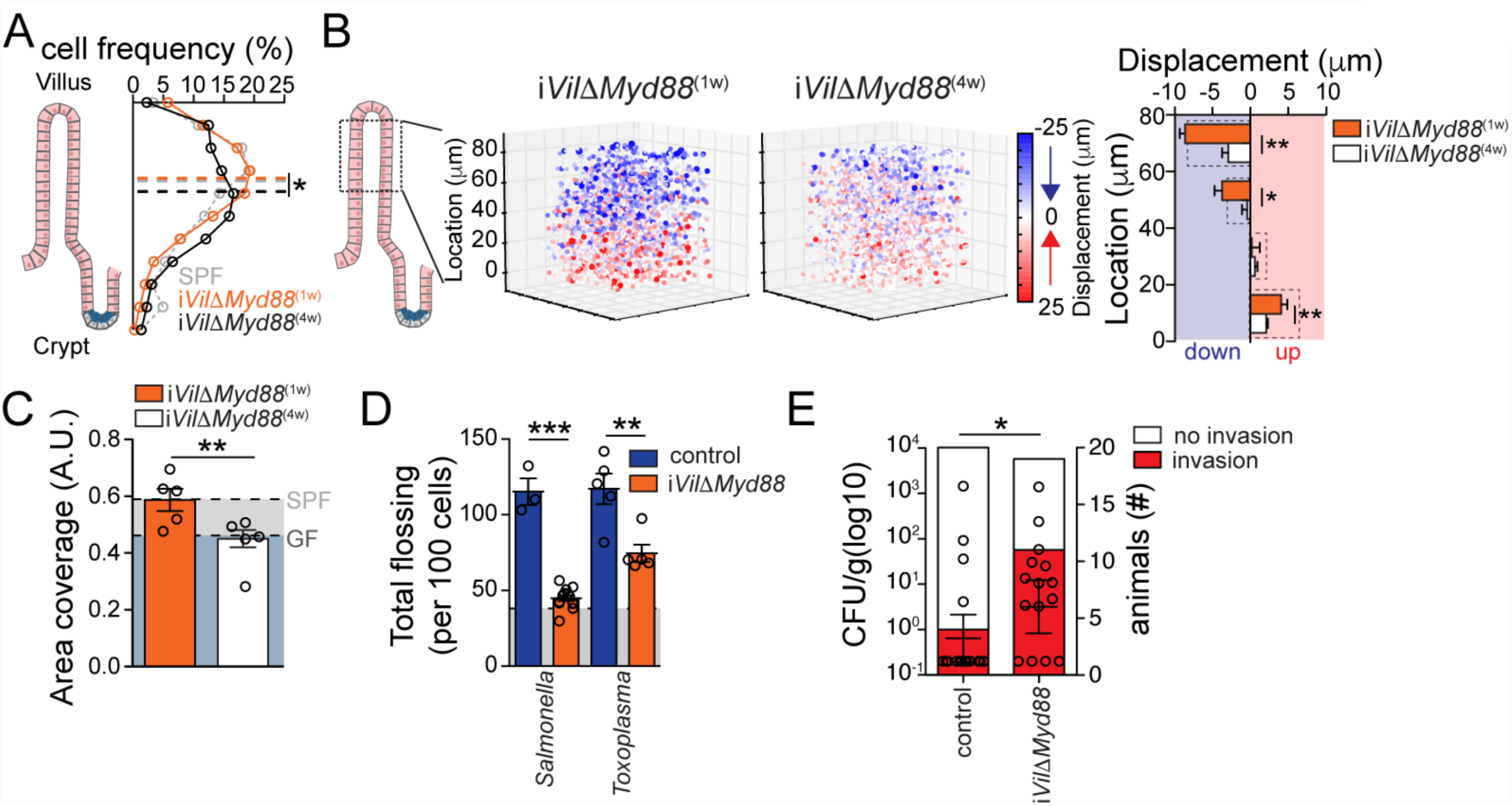
Myd88 expression by ECs modulates IEL dynamic response to intestinal microbes. (A) Frequency of TCRγδ cell distribution along the cleared ileum villi of mice 1 wk or 4 wks after tamoxifen treatment of i*Vil*Δ*Myd88*-TCRγδ^GFP^ mice. Dashed line indicates mean of median positions for 4 mice/group. * = p<0.05 with two-tailed Students *t* test. (B) IVM visualization and quantification of TCRγδ vertical (*Z*) displacement. Pooled imaging data from i*Vil*Δ*Myd88*-TCRγδ^GFP^ mice 1 or 4 weeks after tamoxifen treatment is shown. Panels show starting position and mean vertical displacement over time for each individual cell within each movie. Color density and size indicate degree of *Z* displacement down- (blue) or upwards (red). Graph shows mean *Z* displacement per anatomical villus region as indicated. N= at least 4 mice/group in 3 independent experiments. = p<0.05, ** = p<0.01 with two-tailed Students *t* test. (C) Quantification of unique area covered/IEL/h. Means and SEM shown, each dot = 1 mouse. Light grey shading = SPF value, dark grey shading = GF value (Fig. 1). ** = p<0.01 with two-tailed Students *t* test. (see Movies S4A, B). (D) Tamoxifen-treated (1 wk) i*Vil*Δ*Myd88*-TCRγδ^GFP^ and Cre^−^ littermate control mice were infected with *S.* Typhimurium 18h or *Toxoplasma gondii* (25 cysts) 6h prior to IVM. Unbiased computational quantification of flossing movements. Light grey shading = SPF naïve value (Fig. 1). ** = p<0.01, *** = p<0.001 with two-tailed Students *t* test (see Movies S4C-F). (E) *S.* Typhimurium CFU/g of liver tissue and absolute # of tamoxifen-treated i*Vil*Δ*Myd88*-TCRγδ^GFP^ and Cre^−^ littermate control mice with liver invasion (right axis) 24h after *S.* Typhimurium infection. * = p<0.05, Fisher’s Exact test (right axis), Mann-Whitney *u* test (left axis).

Next, we examined whether γδ IEL behavioral responses to enteric pathogens also require EC-specific Myd88 expression, as suggested by the transcriptional analyses. In contrast to γδ IELs from tamoxifen-treated Cre^-^TCRγδ^GFP^ littermate control mice, upon infection with *Salmonella* or *Toxoplasma* IELs from i*Vil*Δ*Myd88-*TCRγδ^GFP^ mice showed flossing behavior similar to that of naïve animals (Movie S4C, D; Figure 4D). While modulation of γδ IEL behavior was dependent on EC Myd88 expression, we did not detect a role for TCRγδ-ligand interactions, since treatment of TCRγδ^GFP^ mice with anti-TCRγδ blocking antibodies did not prevent increased flossing upon infection (Figure S4A, B). Finally, in a similar fashion to that previously described for total TCRγδ-deficient mice (Ismail et al., 2011), we observed an increased incidence of early invasion of *Salmonella* in tamoxifen-treated i*Vil*Δ*Myd88*, but not in littermate control mice (Figure 4E). The above data demonstrate that ECs are responsible for regulating γδ IEL behavior in both the steady state and during infection in a Myd88-dependent and TCR-independent manner.

### Metabolic regulation of γδ IEL behavior

Metabolic re-programming is essential for T cell clonal expansion and effector function (Buck et al., 2015; Ho et al., 2015; Kidani et al., 2013), although its role in terminally-differentiated (and non-proliferative) tissue-resident T cells is less understood. Because the above transcriptome analyses suggested metabolic changes in γδ IELs in response to infection, we analyzed their energy utilization pathways using extracellular flux analysis (EFA) of primary cells. We observed a rapid increase in extra-cellular acidification rate (ECAR) in sorted γδ IELs from *Salmonella*-infected mice when compared to γδ IELs isolated from naïve control mice, indicating enhanced glycolytic activity in these cells (Figure 5A, B). In addition to its role in modulating IEL behavior, the transcriptome analyses suggested that Myd88 expression by ECs could also influence IEL metabolic programs following infection. To investigate this possibility, we performed EFA on γδ IELs isolated from naïve i*Vil*Δ*Myd88* mice and i*Vil*Δ*Myd88* mice after *Salmonella* infection. Deletion of *Myd88* in ECs resulted in a completely abrogated glycolytic response (ECAR upregulation) by γδ IELs upon *Salmonella* infection, indicating that microbial sensing by ECs is required to regulate both IEL energy utilization as well as their dynamic behavior (Figure 5C).

**Figure 5.**
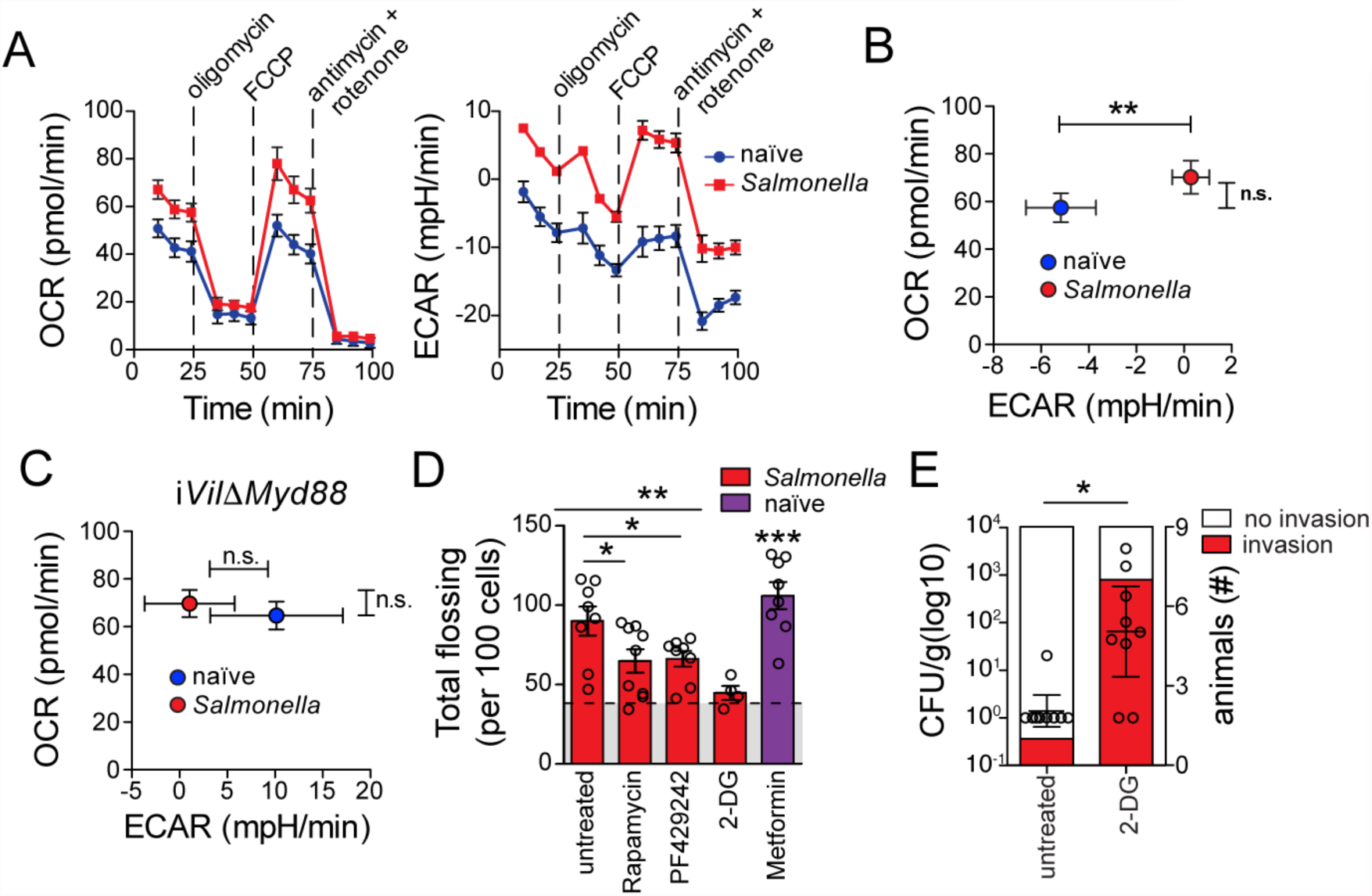
Myd88 expression by ECs modulates IEL metabolic response to intestinal microbes. (A-C) Mitochondrial Stress Test (Seahorse) on sorted TCRγδ IELs isolated from *S.* Typhimurium-infected (18h) or naïve wild-type (A, B) or tamoxifen-treated i*Vil*Δ*Myd88* mice (C). Oxygen Consumption Rate (OCR) and Extracellular Acidification Rate (ECAR) were quantified. N = at least 5 mice/group in 3 independent experiments. ** = p<0.01 with two-tailed Students *t* test. (D) Unbiased computational quantification of flossing movements visualized by IVM after *S.* Typhimurium infection (18h, red) or naïve (purple), and treatment with indicated drugs (see Movies 3A-E). Means and SEM shown, each dot= 1 movie, N= 4-5 mice/group in 3 independent experiments. * = p<0.05, ** = p<0.01, *** = p<0.001 with two-tailed Students *t* test. (E) *S.* Typhimurium CFU/g of liver tissue and absolute # of animals with liver invasion (right axis) 24h after infection in 2-DG treated or control wild-type mice. * = p < 0.05, Fisher’s Exact test (right axis), Mann-Whitney *u* test (left axis).

The gene set enrichment analyses shown in Figure 3 suggest that the mTOR pathway may be involved in the increased glycolytic response observed in γδ IELs post-infection, and this is similar to what has been described for activated effector T cells (Chang et al., 2013; Kidani et al., 2013).

To address whether γδ IEL metabolic changes were coupled with changes in movement dynamics, we administered rapamycin, an inhibitor of mTOR, or PF429242, an inhibitor of SREBP activity downstream of mTOR (Saxton and Sabatini, 2017), to TCRγδ^GFP^ mice prior to *Salmonella* infection. Inhibition of either mTOR- or SREBP-dependent glycolytic responses significantly suppressed IEL flossing behavior post-infection, although speed and vertical displacement patterns along the villi were maintained (Movie S5A-C; Figure 5D, Figure S5A-C). Furthermore, inhibition of mTOR prevented infection-induced flossing hotspots at lower villi regions while it induced hotspots at tip regions (Figure S5C). These effects were also observed by directly blocking glycolysis through the administration of 2-deoxy-glucose (2-DG), a non-metabolizable glucose analog that competes with glucose for hexokinase (Movie S5D; Figure 5D, Figure S5A-C). Conversely, *in vivo* administration of metformin, an mTOR-independent activator of glycolysis, to naïve mice was sufficient to recapitulate γδ IEL dynamic changes observed in response to infection (Movie S5E; Figure 5D, Figure S4A-C). These data suggest that changes in γδ IEL displacement depend on the rapid generation of cytosolic ATP that is achieved by glycolysis. Metformin administration induced increased overall flossing hotspots, although they accumulated throughout the villi, contrasting with the behavior observed after enteric infection (Figure S5C). These data reinforce the possibility that this behavior might be infection site- or epithelial-damage specific. Finally, we determined whether these glycolysis-dependent γδ IEL behavioral changes were linked to early pathogen invasion. While metformin administration did not rescue the increased susceptibility to early *Salmonella* invasion observed in tamoxifen-treated i*Vil*Δ*Myd88* mice, 2-DG treatment of wild-type animals resulted in enhanced bacterial invasion (Figure S5D, Figure 5E). This data suggests that increased glycolytic activity by γδ IELs and their corresponding behavioral response is required but not sufficient for early resistance to infections. Additionally, the above results point to role for pathogen-induced IEL inter-epithelial hotspots in response to luminal insults. Overall, these results identify a coordinated tissue response to microbial cues, in which ECs modulate T cell function in the intestine by directing a metabolic switch coupled with altered cell dynamics and immune function to coordinate immune surveillance of the epithelium.

## Discussion

A large fraction of T cells resides in the intestinal epithelium, thus in close contact with an abundance of diverse microbes, dietary antigens and metabolites; yet how these IELs respond to luminal perturbations is not well understood (Cheroutre et al., 2011; Vantourout and Hayday, 2013). Furthermore, despite the advances in high-resolution *in situ* imaging tools in the past 15 years (Germain et al., 2012), very little understanding of how immune cells dynamically adapt to tissue-specific cues in mucosal tissues has been gained (Edelblum et al., 2012; Edelblum et al., 2015; Farache et al., 2013; McDole et al., 2012; Sujino et al., 2016; Wang et al., 2014). The studies presented here uncover a new layer of tissue regulation of hematopoietic cells. We defined an essential role for microbial sensing by ECs in the regulation of T cell movement dynamics and energy utilization.

First, using live imaging and high-resolution microscopy, we provide several lines of evidence that a major IEL population, γδ T cells, dynamically responds to microbial cues. Although previous studies have demonstrated that “natural” IELs do not require commensal bacteria to populate the intestinal epithelium (Bandeira et al., 1990; Latthe et al., 1994; Staton et al., 2006), we observed that γδ IEL distribution along the small intestine, positioning in the villi, motility above the basement membrane and villus-crypt bidirectional displacement are largely dependent on stimulation by the microbiota and enteric pathogens. It is possible that IELs physiologically sense EC proliferation, since our estimated IEL vertical downward displacement nearly offsets the EC growth rate described in SPF mice (approximately 6 μm/hour) (Park et al., 2016). Likewise, this coordination seems to be directed by gut microbiota, since both γδ IEL vertical displacement and epithelial cell turnover are reduced in mice devoid of microbiota (Park et al., 2016). The microbiota-dependent preferential localization of γδ IELs in the upper third of villi also corresponds to the increased microbial concentration reported in that region, which has lower concentrations of anti-microbial peptides (AMPs) produced by Paneth cells in the crypts (Clevers and Bevins, 2013; Hooper and Macpherson, 2010). Therefore, our data suggest that microbial density, not only along the gastrointestinal tract but also around individual villi, may serve as a major driver of γδ IEL localization and surveillance of the epithelium. Rather than direct interaction between γδ IELs and luminal bacteria, the observed microbial effects on γδ IELs were dependent on sensing and instruction by tissue cells.

Several pro- and anti-inflammatory mechanisms depend on bidirectional interactions between hematopoietic and non-hematopoietic tissue cells. For instance, tissue-resident memory T cell precursors are constantly recruited to the intestinal epithelium in the absence of infection and to additional mucosal sites and the skin during infection or inflammatory responses (Laidlaw et al., 2014; Mackay et al., 2016; Mackay et al., 2013; Masopust et al., 2001; Schenkel et al., 2014; Shin and Iwasaki, 2013). Additionally, γδ IELs have been linked to wound repair and maintenance of the gut and skin epithelial barrier (Boismenu and Havran, 1994; Chen et al., 2002; Yang et al., 2004). We found that in conditions of depleted microbiota, reduced γδ IEL vertical villus-crypt displacement was directly correlated with reduced EC proliferation raising the possibility that physiological signals associated with EC growth serve to fine-tune T cell dynamics in the tissue. The fact that γδ IELs located at the tips of the villi showed a strong downward movement trend (and vice-versa) suggests the existence of repulsive signals at the extremities and/or attraction signals at the middle of the villus. Based on our parallel EC-IEL transcriptome analyses, we speculate that one of those signals may involve the β-Catenin/Wnt pathway, which maintains the self-renewal capacity of epithelial stem cells (Clevers et al., 2014; Karin and Clevers, 2016).

We observed that enteric infections resulted in fast and well-defined behavioral changes in γδ IELs, including reduced vertical displacement, altered villus positioning and heightened inter-epithelial flossing movement. This infection-induced flossing movement preferentially occurred in specific hotspot areas after either *Toxoplasma* or *Salmonella* infection. Data obtained using the *Salmonella* mutant *invA*, as well as in mice 3 days after *Toxoplasma* infection, suggest that these behavioral changes require continued presence of the pathogen. We speculate that flossing hotspots and altered γδ IEL vertical displacement could be related to a specific response to tissue damage or high concentrations of enteric pathogens in particular regions of villi. This possibility is consistent with an earlier report using live imaging of intestine bathed in *Salmonella-*containing medium, although in this case, the authors observed large number of TCRγδ cells in the lamina propria prior to infection (Edelblum et al., 2015). In the present study and our previous work, we find that only a small fraction of TCRγδ (< 10%) is present in the LP (Sujino et al., 2016). Supporting an active role intestinal epithelial barrier function, previous studies have demonstrated that TCRγδ-deficient mice are highly susceptible to both of these pathogens, and these mice show early pathogen invasion to peripheral organs and systemic sites (Dalton et al., 2006; Ismail et al., 2011). γδ IELs have also previously been shown to play a role in stress-surveillance (Hayday, 2009). Imaging studies of γδ IELs in the murine skin supported this concept, however skin IELs largely display a dendritic cell-type morphology with limited migration (Chodaczek et al., 2012), which contrasts with the intestinal IEL lymphoid morphology and mobile phenotype (Edelblum et al., 2012; Sujino et al., 2016).

Although the exact functional relevance of T cell inter-epithelial movement triggered by infection remains unknown, studies using TCRγδ-deficient mice in models of infection point to a role for γδ IELs in the maintenance of intestinal epithelial tight junctions during acute phases of infection (Dalton et al., 2006; Edelblum et al., 2015; Ismail et al., 2011). The concomitant changes in expression of genes in pathways related to cell adhesion in ECs and γδ IELs post-infection reinforces this possibility. Additionally, production of cytokines such as IL-15 by ECs contributes to IEL cytotoxic activity during intracellular infections, although exaggerated IL-15 production during inflammation can trigger IEL-mediated destruction of the epithelial barrier and disorders such as coeliac disease (DePaolo et al., 2011; Jabri and Abadie, 2015; Meresse et al., 2004; Meresse et al., 2006; Tang et al., 2009). Of note, flossing behavior would provide γδ IELs enhanced access and proximity to pathogens during infection. Our observed transcriptional changes in genes associated with NK and cytotoxic activity in γδ IELs upon infection, supported by previous accounts showing that these cells are able to upregulate anti-microbial molecules (Ismail et al., 2011), raise the possibility of direct anti-bacterial or -protozoan activity by γδ IELs. Previous studies have demonstrated several roles for EC pattern recognition receptor signaling in the maintenance of host-microbiota equilibrium, including in pathogen-defense, epithelial cell growth and repair, and conditioning of innate immune cells (Dessein et al., 2009; Rakoff-Nahoum et al., 2004; Slack et al., 2009; Staton et al., 2006; Vaishnava et al., 2008; Zaph et al., 2007). We described that specific ablation of the gene encoding the adapter protein Myd88 on ECs was sufficient to recapitulate the γδ IEL behavioral phenotype and tissue distribution observed in mice devoid of microbiota. γδ IEL responses to enteric pathogens, including flossing movements, were also impaired in mice lacking Myd88 on ECs. Furthermore, EC microbial pattern-sensing was found to be required for a γδ IEL metabolic switch in response to pathogens. A requirement for increased glycolysis has been demonstrated in T cells transitioning from naïve and memory states into an effector state (Chang et al., 2013; Macintyre et al., 2014; Vander Heiden et al., 2009). However, an increased glycolytic rate in effector T cells is normally linked with TCR activation and cell proliferation (Buck et al., 2015), neither of which were observed in γδ IELs in response to infection, resembling what has been proposed for innate immune cells (Krawczyk et al., 2010). In the absence of infection, induction of glycolysis was sufficient to trigger an array of γδ IEL behavioral changes normally observed after enteric infections, while inhibition of glycolysis prevented these infection-induced γδ IEL dynamic changes. While the significance of this concomitant EC-dependent modulation of γδ IEL metabolism and behavior to gut physiology remains to be determined, targeting either EC pathogen sensing or glycolysis led to increased susceptibility to *Salmonella* invasion, closely mirroring the phenotype of mice deficient in γδ T cells (Ismail et al., 2011). Therefore, ECs appear to synchronize γδ IEL surveillance to control enteric pathogen translocation through the epithelial layer. The potential role for glucose utilization in T cell surveillance of the gut epithelium contrasts to the beneficial effects of fasting metabolism in preventing neuronal dysfunction during sepsis (Wang et al., 2016), raising the possibility that tissue-specific physiology and infection routes, in addition to particular pathogen characteristics, influence specific metabolic requirements for pathogen resistance and tissue tolerance.

In conclusion, we have described an intricate and dynamic relationship between intestinal epithelial cells and intra-epithelial lymphocytes, where EC sensing of microbes determines γδ IEL tissue distribution, migration and scanning patterns as well as energy utilization. This study thus provides support for an EC-dependent, coordinated tissue-immune cell response to environmental insults.

## Author Contributions

D.M. conceived; D.M and B.S.R. supervised this study; D.vK, B.S.R., V.P. and D.M. designed experiments; D.vK., B.S.R. and V.P. performed and analyzed experiments; J.F. and G.V. helped setting up and performed initial IVM experiments. D.vK., B.S.R., V.P., J.F. and G.V. prepared figures and helped with manuscript preparation; D.M. wrote the paper.

## Acknowledgments

We are indebted to N. Thomas and K. Gordon for sorting cells, to Thiago Kikuchi Oliveira for assistance with RNA sequencing analysis and other members of the Nussenzweig lab and The Rockefeller University employees for continuous assistance. We thank Sylvie Robine (Institut Curie) for providing *Vil*^CreER^ mice and J. Cross (Memorial Sloan Kettering Cancer Center) for support with the metabolic studies. We thank Guy Shakhar (Weizmann Institute) for discussions regarding initial imaging experiments. We especially thank A. Rogoz for outstanding technical support. We thank members of our laboratory, particularly A. Lockhart, and D. Esterhazy, for discussions and critical reading and editing of the manuscript. This work was supported by Leona M. and Harry B. Helmsley Charitable Trust (B.R., D.HvK., V.P., D.M.), Alexandre Suerman Stipend, Royal Netherlands Academy of Sciences and the Prince Bernhard Cultural Foundation (D.HvK.), the Crohn’s & Colitis Foundation of America (B.R., D.M.), and National Institutes of Health NIH R01 DK093674 and R01 DK113375 grants (D.M.). The Rockefeller University Bio-Imaging Resource Center is supported by the Empire State Stem Cell Fund through NYSDOH C023046.

## Supplemental Figures

**Supplemental Figure 1. Influence of microbiota on the distribution of IEL subpopulations in the small intestine**

(A) Flow cytometry analysis of frequency and total cell count of IEL populations in the duodenum (D), Jejunum (J) and ileum (I) of wild type specific pathogen-free (SPF) and germ-free (GF) mice. Cell number was adjusted by the size of each region in centimeter. IEL populations were gated as: TCRγδ^+^ CD8α^+^; TCRαβ^+^CD8αα^+^; TCRαβ^+^CD8αβ^+^; TCRαβ^+^CD4^+^CD8β^-^; TCRαβ^+^CD4^+^CD8β^−^CD8α^+^ (all among CD45^+^ cells). (B, C) Representative images of cleared (*FocusClear*™) jejunum and ileum villi of SPF and GF TCRγδ^GFP^ mice. In green (GFP), TCRγδ^+^ cells and in blue (Hoechst), EC nuclei. (D) Means and SEM of TCRγδ^GFP^ cell speed are shown, each dot= 1 movie, N= 4-5 mice/group in 3 independent experiments. ** = p<0.01 with two-tailed Students *t* test. **Related to Figure 1.**

**Supplemental Figure 2. Influence of enteric infections on IEL behavior**

(A) Quantification of unique area covered/IEL/hour (measured with IVM) after infection with *Salmonella* and *Toxoplasma* at indicated time points. (B) Unbiased computational quantification of flossing movements after infection with *Salmonella* or *Toxoplasma* at indicated time points and visualized by IVM (see Movies S3A-C). % of flossing movements occurring in a “hotspot” area is shown for each villus region, N= 4-5 mice/group in 3 independent experiments. (C) Representative images of cleared (*FocusClear*™) duodenum, jejunum and ileum villi of *S.* Typhimurium*-*infected TCRγδ^GFP^ mice after 2h or 18h of infection. In green (GFP), TCRγδ^+^ cells and in blue (Hoechst), EC nuclei. (D) Means and SEM of TCRγδ^GFP^ cell speed are shown after 6h infection with *Toxoplasma* (25 cysts), each dot= 1 movie, N= 4-5 mice/group in 3 independent experiments. **Related to Figure 2.**

**Supplemental Figure 3. Myd88 expression by ECs is required for a coordinated epithelial transcriptional response to microbes**

Sorted TCRγδ IELs (CD45^+^EpCAM^-^TCRαβ^-^TCRγδ^+^CD8α^+^) (A) and ECs (CD45^-^EpCAM^+^) (B) were isolated from naïve wild-type mice or from tamoxifen-treated *S.* Typhimurium-infected i*Vil*Δ*Myd88* and Cre^−^ littermate control mice. Individual gene expression (row normalized) is shown for genes involved in indicated GO Biological Processes for (A) TCRγδ IELs and (B) ECs. Only genes with minimal expression exceeding 0.5 RPKM (top 30 %) are shown. **Related to Figure 3.**

**Supplemental Figure 4.**

(A). *Nur77*^GFP^ mice were i.p. injected with 400 μg of anti-murine TCRγδ antibody (clone UC7-13D5) or isotype control in 100 μl PBS at days −1, 0 before cell harvest. IELs were *in vitro* stimulated with 5 μg of plate-bounded α-CD3 and α-TCRγδ (clone GL-3) for 3 h at 37°C and stained for FACS analysis. (B) Unbiased computational quantification of inter-epithelial cell movements (flossing) in wildtype TCRγδ^GFP^ mice after with *S.* Typhimurium infection (18h). Anti-TCRγδ monoclonal antibody or isotype control was administrated intraperitoneally at -1 and 0 days prior to infection as in C. Each dot = 1 mouse. **Related to Figure 4.**

**Supplemental Figure 5. Myd88 expression by ECs modulates IEL metabolic response to intestinal microbes**

(A) Means and SEM of TCRγδ^GFP^ cell speed are shown (measured with IVM) after *S.* Typhimurium infection (18h, red) or naïve (purple), and treatment with indicated drugs. Each dot= 1 movie. N = at least 4 mice/group in 3 independent experiments. Grey shaded area = SPF naïve value (Fig. 1). *** = p<0.001 (vs. SPF naïve) with two-tailed Students *t* test. (B) Graph shows mean *Z* displacement per anatomical villus region (measured with IVM) as indicated after *S.* Typhimurium infection (18h) or naïve, and treatment with indicated drugs. Grey shading indicates naïve wildtype level of movement as measured in Figure 1. N = at least 4 mice/group in 3 independent experiments. = p<0.05, with two-tailed Students *t* test. (C) Unbiased computational quantification of flossing movements in naïve mice or after infection with *Salmonella* (18h) and treatment with indicated drugs visualized by IVM (see Movies S5A-E). % of flossing movements occurring in a “hotspot” area is shown for each villus region. N= 4-5 mice/group in 3 independent experiments. (D) *S.* Typhimurium CFU/g of liver tissue and absolute # of animals with liver invasion (right axis) 24h after infection in Tamoxifen-treated (1 wk.) i*Vil*Δ*Myd88* mice, with or without metformin treatment. **Related to Figure 5.**

## Methods

### Contact for Reagent and Resource Sharing

Further information and requests for resources and reagents should be directed to and will be fulfilled by Daniel Mucida.

### Experimental Model

#### Mice

C57BL/6 (000664), *Nur77*^GFP^ (016617) and TCRγδ^GFP^ (016941), mice were purchased from the Jackson Laboratories and maintained in our facilities. *Villin*^Cre-ERT2^ mice were generated by Sylvie Robine (Institut Curie) and provided by David Artis (Cornell Univ.); *Myd88^f/f^* mice were provided by Michel Nussenzweig (Rockefeller Univ.). These lines were interbred in our facilities to obtain the final strains described in the text. Mice were maintained at The Rockefeller University animal facilities under specific pathogen-free (SPF) or germ-free (GF) conditions. C57BL/6 GF mice were originally obtained from Sarkis Mazmanian (Caltech) and maintained in our facility. TCRγδ^GFP^ GF mice were rederived, bred and maintained in our GF facilities. GF status was confirmed by plating feces as well as by qPCR analysis (16S rRNA). Mice were used at 7-12 weeks of age for all experiments except when otherwise indicated. Littermates of the same sex were randomly assigned to experimental groups. Both female and male mice were used for experiments, no notable sex-dependent differences were found for the reported experiments. Additionally, we infected and analyzed mice at the same time of day, maintaining a similar time of analysis between experiments. Animal care and experimentation were consistent with the NIH guidelines and were approved by the Institutional Animal Care and Use Committee at The Rockefeller University.

#### Microorganisms

*Salmonella enterica* serovar Typhimurium (SL1344) and its mutant *invA* were used for infection experiments and cultured prior to infection as described below. *Toxoplasma gondii* me49 were generously provided by Ricardo T. Gazzinelli (U. Mass.) were maintained in our lab by periodically infecting C57BL/6 mice with 5 cysts administrated intraperitoneally (i.p.)- cysts were removed from their brain 30 days after i.p. infection.

### Method Details

#### Antibodies and flow cytometry analysis

Fluorescent-dye-conjugated antibodies were purchased from BD-Pharmingen (anti-CD4, 550954; 557495; anti-CD45R, 557683;) or eBioscience (anti-CD8α 56-0081; anti-TCR-αβ 47-5961; anti-TCRγδ, 46-5711; anti-CD8β, 46-0083;). Flow cytometry data were acquired on an LSR-II flow cytometer (Becton Dickinson) and analyzed using FlowJo software (Tree Star). For TCR blocking of TCRγδ cells 400 μg of anti-TCRγδ monoclonal antibody (UC7-13D5, ATCC CRL-1989) was administrated i.p. at -1 and 0 days prior to infection. Total cell count was performed using CountBright absolute counting beads (Invitrogen) as instructed by the manufacturer.

#### RNA-sequencing

RNA was isolated from samples using the RNAeasy kit (Qiagen, USA) according to the instructions provided by the manufacturer. RNA libraries from biological replicates were prepared using the SMART-Seq™ v4 Ultra™ Low Input RNA (ClonTech Labs) and/or TruSeq^®^ Stranded mRNA Sample Preparation and sequenced using 75 base pair single end high-output reading on a NextSeq 500 instrument (Illumina). The reads were aligned using the STAR version 2.3.0 software that permits unique alignments to Mouse Ensembl genes. Differential expression was determined by use of the Cufflinks software with default settings. Gene Set Enrichment Analysis (GSEA PreRanked, Broad Institute)(Subramanian et al., 2005) of differentially expressed genes was performed with default settings to identify enrichment of curated gene sets derived from the Gene Ontology Biological Processes pathway database (www.geneontology.org). Gene ontology analysis of individual genes and pre-selected groups of genes (as in Figure 3E, F) was performed using the PANTHER classification system (www.pantherdb.org) (Mi et al., 2013) and the Gene Ontology consortium (www.geneontology.org). False Discovery correction was applied as detailed in the figure legends.

#### Preparation of intraepithelial lymphocytes

Intraepithelial were isolated as previously described (Mucida et al., 2007). Briefly, small and large intestines were removed and placed in chilled HBSS media (Gibco) containing 2% FCS. The intestines were carefully cleaned from the mesentery and flushed of fecal content.

Intestines were opened longitudinally and then cut into 1 cm pieces. The intestinal tissue was transferred to a 50 ml Falcon tubes containing 25 ml of cold HBSS complemented with 2% FCS and 5 mM EDTA and shaken (2x) at 230 rpm for 20 min at 37°C. The tissue suspension was passed through a stainless steel sieve into 50-ml conical tubes and the cells were pelleted by centrifugation at 1200 rpm for 10 min at 4°C. The cell pellet was resuspended in complete HBSS, layered over a discontinuous 40/70% Percoll gradient, and centrifuged at 2000 rpm for 30 min. Cells from the 40/70% interface were collected, washed and resuspended in complete RPMI media. These purified cells constituted the intraepithelial lymphocyte (IEL) population.

#### Multiphoton microscopy

TCRγδ^GFP^ reporter mice were anesthetized with i.p. injection of 20 μL/g of 2.5% Avertin before surgery for intravital imaging. Anesthesia was maintained by continuous administration 1% of isofluorane and 1L per minute of oxygen mixture while imaging was performed. Mice were injected with Hoechst dye (blue) for visualization of epithelial cell nuclei. 10 min following induction of anesthesia, mice were placed on a custom platform heated to 37°C. Upon loss of recoil to paw compression, a small incision was made in the abdomen. The ileum entrance to the caecum was located and a loop of ileum was exposed and placed onto a raised block of thermal paste covered with a wetted wipe. A coverslip was placed on top of the loop to immobilize the intestine. The platform was then transferred to the FV1000MPE Twin upright multiphoton system (Olympus) heated stage. Time-lapse was +/- 30sec with a total acquisition time of 30 min. A complete Z-stack (80 μm) of several ileum villi was made during each acquisition. *Imaris* (Bitplane AG) software was used for cell identification and tracking, using the “*Spots”* auto-regressive tracking algorithm as described by the manufacturer. The scoring of IEL behavior (flossing, speed, *z*-movement, location) was entirely computational and unbiased, using Python (Enthought Canopy) integrated with Microsoft Excel and with the application of standard scientific algorithm packages. “Flossing” movements are identified using extensively verified parameters, based on their unique properties (e.g. sequence of specific sharply angled movements). See below for additional detail on quantification and statistical analysis of imaging data.

#### Metabolism assays

Extracellular acidification (ECAR) and oxygen consumption rate (OCR) were measured for sorted *ex vivo* TCRγδ cells using a 96-well Seahorse XFe96 Analyzer (Agilent / Seahorse Bioscience) in the Cell Metabolism Laboratory of the Donald B. and Catherine C. Marron Cancer Metabolism Center at Memorial Sloan Kettering Cancer Center (MSKCC) using the Seahorse XF Mito Stress Kit (Agilent) according to the manufacturer’s instructions with titrated concentrations of mitochondrial inhibitors and measures for lymphocyte adhesion. Briefly, sorted T cells were adhered to assay plates coated with 25 μg/mL Cell-Tak (Corning) in 0.1 M NaHCO_3_ at pH 8.0 and incubated in the absence of CO_2_ in assay medium (non-buffered DMEM containing 10 mM glucose, 2 mM L-glutamine, and 1 mM sodium pyruvate) for 45 min before the assay. ECAR and OCR were measured under basal conditions and in response to sequential addition of 2 μM oligomycin, 2 μM carbonyl cyanide-4 (trifluoromethoxy) phenylhydrazone (FCCP) and 0.5 μM rotenone + 0.5 μM antimycin A at indicated time-points.

#### Whole Mount Immunofluorescence (iDISCO and *FocusClear™*)

The iDISCO protocol was followed as detailed on the continuously updated website htpp://idisco.info. Briefly, mice were sacrificed by cervical dislocation and the small intestine was removed and placed in HBSS Mg^2+^Ca^2+^(Gibco) + 5% FCS. The intestine was cut open longitudinally and the luminal contents washed away in complete media with 1 mM DTT (Sigma-Aldrich). The tissue was pinned down in a plate coated with Sylgard and then fixed for O/N with 4% PFA at gentle agitation. After washing and methanol dehydration steps, whole mount samples were then permeabilized first in 0.2% Triton X-100 for 2 hours followed by 0.2% Triton X-100/20%DMSO/0.3MGlycine for 1-2 days at 37°C with gentle agitation. After washing in 1X DPBS with 0.2% Tween-20 and Heparin (100 μg/ml), the samples were blocked for 1-2 days in 1X DPBS with 0.2% Triton X-100/10%DMSO/6%Donkey Serum for 1-2 days at 37°C with gentle agitation. Antibodies were added to the blocking buffer at appropriate concentrations and incubated 1-2 days at 37°C. After primary incubation the tissue was washed 5 – 10 X in 1X DPBS with 0.2%Tween-20 and Heparin (100 μg/ml), and then incubated in blocking buffer with secondary antibody at concentrations within the primary antibody range. Samples were again washed 5 – 10 X in 1X DPBS with 0.2%Tween-20 and Heparin (100 μg/ml), mounted in agarose and cleared using dichloromethane followed by benzyl ether. Images were taken on an Ultramicroscope (LaVision BioTec) with light sheet illumination and adjusted post-hoc in Imaris. The following unconjugated primary antibodies were used to stain the intestine: Aves Labs (anti-GFP chicken polyclonal), Invitrogen (anti-ZO-1 / TJP1 rabbit polyclonal). *FocusClear™* tissue clearing was performed according to manufacturer’s (Celexplorer Labs. Co.) instructions. Briefly, TCRγδ^GFP^ reporter mice were injected with Hoechst dye (blue) for visualization of epithelial cell nuclei. After 15-30 minutes, mice were sacrificed by cervical dislocation and segments of the small intestine were removed, washed (intact) and fixed in 4% PFA at gentle agitation for 2 h at RT. After fixation, samples were washed in 1X DPBS and placed in *FocusClear™* solution for approximately 15 min. at room temperature. Once visual confirmation of clearance was obtained, samples were mounted in 3D printed slides with *MountClear™*, sealed and imaged using an inverted LSM 880 NLO laser scanning confocal and multiphoton microscope (Zeiss). As this protocol maintains native GFP and introduced Hoechst fluorescence, no antibody staining was necessary.

#### Infections

Mice were orally inoculated with 10^8^ CFU of *Salmonella* Typhimurium (SL1344) or mutant *invA*. Imaging and or gene expression analysis was performed 2 - 18 hours after inoculation as indicated. Briefly, a single aliquot of *Salmonella* was grown in 3 ml of LB overnight at 37°C with agitation, and then the bacteria were sub-cultured (1:30) into 3 ml of LB for 3.5h at 37°C with agitation. The bacteria were next diluted to final concentration in 1 ml of DPBS. Bacteria were inoculated by gavage into recipient mice in a total volume of 100μl.

For imaging experiments with *Toxoplasma gondii* infection, mice were infected by oral gavage with the amount of cysts as indicated. Mice were imaged 6 – 72 hours after inoculation as indicated.

##### *Salmonella* invasion of peripheral organs

Whole livers were removed post mortem (without the gall bladder) and homogenized in PBS with 0.1% Triton X-100 to release intracellular bacteria. Colony-forming units (CFU) in the liver were determined by plating 3 serial dilutions of the tissue suspension on selective agar, *Salmonella*-*Shigella* Agar (BD 211597), and resulting quantities were normalized to organ weight prior to homogenization.

##### Antibiotic Treatment

Mice were given ampicillin (1 mg/ml), vancomycin (0.5 mg/ml), neomycin sulfate (1 mg/ml), and metronidazole (0.5 mg/ml) in drinking water with for 4 weeks. Sucralose-based artificial sweetener (Splenda^®^) was added at 5 g/l to both antibiotics-treated and control mice drinking water. All antibiotics were purchased from Sigma. Microbiota depletion was verified by aerobic and anaerobic culture of intestinal contents.

##### Drug treatments

For SREBP pathway inhibition, animals were i.p. injected every 6-8 hours with 10 mg/kg of SREBP inhibitor (PF429242, Tocris Cat# 3354, 20 mg/ml in DMSO) on day -1, 0 and +1 of *Salmonella* infection. For mTOR inhibition, mice were treated with daily i.p. injections of 500 μM Rapamycin (Sigma Cat# R0395) suspended in 0.9% NaCl and 2% ethanol in 100 μL PBS on day -1, 0 and +1 of *Salmonella* infection. For glycolysis inhibition, animals were treated with i.p. injections of 400 mg/kg of 2-DG (Sigma Cat# D6134) on day -1, day 0 and day +1 of *Salmonella* infection. For glycolysis pathway activation, animals were injected with 250 mg / kg Metformin (USP Cat#136309) in 100 μL PBS daily for 3 days and cells analyzed 2-3 hours after last injection.

### Quantification and Statistical Analysis

#### Statistical approach and sample size calculation

Statistical analysis was performed in GraphPad Prism software. For large datasets preliminary or additional statistics were performed using Python (Enthought Canopy) integrated with Microsoft Excel and with the application of standard scientific algorithm packages (see below for additional details) obtained from SciPy (www.scipy.org) and Enthought. For specific statistical tests used in each experiment as well as definition of center, and dispersion and precision measures please refer to figure legends. Our general approach is described below.

For IVM experiments evaluating IEL behavioral patterns we analyzed the difference between two independent means (two groups) using a two-tailed Student’s *t* test. A level of α=0.05 was used to define significance. Power calculations for IVM behavioral experiments using G*POWER software (Faul et al., 2007), based on the minimal effect size of d = 2.336 (preliminary data) showed a requirement of minimum *n*=5 per group. For tissue clearing experiments, we analyzed the difference between two independent means (reflecting mean position of IELs per sample) per group using a two-tailed students *t* test. Power calculations performed as above showed a sample requirement of *n* = 9 per group (3 independent experiments with 3 mice/condition). For RNA-seq, *n*=3/group was compared, as suggested based on extensive review of statistical power for RNA-seq analysis (Conesa et al., 2016), showing a power level of > 85% with our sequencing approach. False Discovery Rate correction was applied for all analyses. For metabolic analyses analyzed the difference between two independent means using a two-tailed Student’s *t* test. Analysis of our preliminary data showed an effect size of d = 1.31 for mean *ex vivo* ECAR, leading to a power calculation (as above) requiring *n* = 14 per group (3 independent experiments with 5 mice/condition). Tissue invasion by pathogens was compared at different time points with a two-tailed Mann-Whitney *u* test as well as the incidence of invasion (as a categorical variable) per group with Fisher’s Exact test. Power calculations suggested a sample size of n=11 (3 independent experiments with 3-4 mice/group).

#### Computational analysis of multiphoton microscopy

For each individual movie, time-lapse was +/- 30sec with a total acquisition time of 30 min. A complete Z-stack (80 μm) of several ileum villi was made during each acquisition. *Imaris* (Bitplane AG) software was used for cell identification and tracking, using the “*Spots*” auto-regressive tracking algorithm as described by the manufacturer. Tracking of cells and identification of IELs was done using the same algorithm for all samples included and manual verification of correct lymphocyte identification will be performed. After this step, all coordinates are inserted into custom-made, standardized algorithms, which analyze lymphocyte coordinates over time to establish IEL localization as well as movement dynamics. The scoring of IEL behavior (flossing, speed, *z*-movement, location) was entirely computational and unbiased, using Python (Enthought Canopy) integrated with Microsoft Excel and with the application of standard scientific algorithm packages obtained from SciPy (www.scipy.org) and Enthought. Specifically, the following packages were used: NumPy, IPython, Matplotlib and Pandas. FFmpeg (www.ffmpeg.org) was used for producing animations of 3D plots. “Flossing” movements are identified using extensively verified parameters, based on their unique properties (e.g. sequence of specific sharply angled movements), the following algorithm is used:

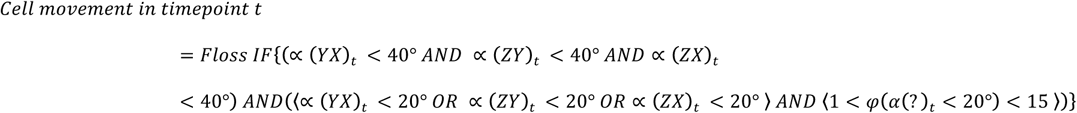

This algorithm defines a flossing movement as a sharply angled movement (< 20 degrees) in 1 dimension, within a defined 3-dimensional space (< 40 degrees in all other dimensions) and with a length of movement between 1 and 15 μm. The accuracy of this analysis was confirmed by visual comparisons and virtual test coordinate sets containing flossing movements.

For analysis of *flossing* hotspots (ie. overlapping *flossing* events), the individual coordinates of each *flossing* experiment in a movie are compared to others, and multiple events per 125 μm^3^ (5 by 5 by 5 μm) volume (the minimum volume an IEL would occupy) scored.

For analysis of vertical (*Z)* movement patterns, mean location in *Z* of each cell per movie was compared to the starting location. Next, the raw starting location with *Z* migration information was plotted in 3 dimensions using *matplotlib* with 3D packages for Python. To analyze differences in *Z* migration patterns between conditions, the net *Z* migration (in μm) was compared for each region of 20 μm (from close to the villus tip to above the crypt, as shown in the graphical representation for each figure). For statistical comparisons, the mean of net *Z* migration per region of several individual animals from independent experiments was compared per condition.

#### Data and Software Availability

Data generated with RNA sequencing was deposited in the NIH GEO database under accession # GSE97184. All software used is available on-line, either freely or from a commercial supplier. Algorithms described above are purely mathematical calculations and can be performed using any software allowing calculations with data matrices. No new software was written for this project.

## Supplemental movie legends

**Supplemental Movie 1 (pertaining to Figure 1) Steady-state localization of intestinal IELs. (Part A)** TCRγδ^GFP^ reporter mice were anesthetized and subsequently perfused with PBS 1X followed by 4% paraformaldehyde (PFA). A small section of the ileum was prepared following the iDISCO protocol as detailed in the methods section. Samples were stained with anti-ZO-1 (red) and anti-GFP (green) antibodies after which imaging was acquired using an Ultramicroscope (LaVision BioTec) with light sheet illumination and adjusted post-hoc in Imaris. The complete Z-stack (200 - 300 μm) of several ileum villi is visible within the field of vision. *Imaris* (Bitplane AG) software was used for data processing using the standard *Spots* algorithm to identify cells compared to background. **(Part B)** TCRγδ^GFP^ reporter mice were anesthetized and subsequently perfused with PBS 1X followed by 4% paraformaldehyde (PFA). A small section of the ileum was prepared for an additional 2 hours of fixation in 4% PFA. Tissue clearing was performed using FocusClear™ (CelExplorar Labs Co.), preserving the native GFP, after which imaging was acquired using an inverted LSM 880 NLO laser scanning confocal and multiphoton microscope (Zeiss). The complete Z-stack (200 - 300 μm) of several ileum villi is visible within the field of vision, with the TCRγδ cells in green (GFP). *Imaris* (Bitplane AG) software was used for data processing using the standard *Spots* algorithm to identify cells compared to background.

**Supplemental Movie 2 (pertaining to Figure 1) Steady-state behavior of intestinal IELs.** SPF (wild type, **Part A**, broad-spectrum antibiotics-treated, **Part C**) and Germ Free (wild type **Part B**, 1 week after re-conventionalization, **Part D**) TCRγδ^GFP^ reporter mice were anesthetized and a small section of the ileum was prepared for *in vivo* imaging, leaving the rest of the bowel intact. Mice were injected with Hoechst dye (blue) for visualization of epithelial cell nuclei. Imaging was acquired using a FV1000MPE Twin upright multiphoton system (Olympus). Timelapse was +/- 30sec with a total acquisition time of 30min. The complete Z-stack (80 μm) of several ileum villi is visible within the field of vision, with the TCRγδ cells in green. A dot indicates a cell is being tracked, with the tracks shown in a time color-coded manner. Tracking was filtered for auto fluorescence. *Imaris* (Bitplane AG) software was used for data processing, with an adapted tracking algorithm. Part B, C and D contain insets of the SPF naïve (Part A) and GF naïve (Part B) movies as indicated for comparison.

**Supplemental Movie 3 (pertaining to Figure 2) Influence of enteric infections on IEL behavior.**

IVM, using the identical image acquisition protocol from Movie S2, is shown for TCRγδ^GFP^ reporter mice after infection with *Salmonella* Typhimurium for 2 hours (**Part A**) or 18 hours (**Part B**), or with *Toxoplasma gondii* (25 cysts for 6 hours, **Part C**). All parts contain insets of Movie S2A (SPF naïve) for comparison.

**Supplemental Movie 4 (pertaining to Figure 4) Myd88 expression by ECs modulates IEL dynamic response to intestinal microbes.**

IVM, using the identical image acquisition protocol from Movie S2, is shown for steady state i*Vil*Δ*Myd88*-TCRγδ^GFP^ mice 1 wk. (**Part A**) or 4 wks. (**Part B**) after tamoxifen treatment (with Part A as inset for comparison). (**Part C**) Tamoxifen-treated i*Vil*Δ*Myd88*-TCRγδ^GFP^ were infected with *S.* Typhimurium 18h or (**Part D**) *Toxoplasma gondii* (25 cysts) 6h prior to imaging as above. Part C and D contain insets of infected Cre – control imaging as indicated for comparison.

**Supplemental Movie 5 (pertaining to Figure 5) The IEL metabolic response to intestinal microbes modulates their dynamic response.**

IVM, using the identical image acquisition protocol from Movie S2, is shown for TCRγδ^GFP^ reporter mice after *S.* Typhimurium infection (18h, **Part A - D**) or naïve (**Part E**), and treatment with indicated drugs. Parts B, C and D contain inset of Part A for comparison. Part E contains inset of Movie S2A (SPF naïve) for comparison.

